# Bayesian phylogenetic inference accounting for recombination in whole bacterial genomes

**DOI:** 10.64898/2026.11.04.764181

**Authors:** Xavier Didelot, Daniel Falush, Daniel J Wilson

**Affiliations:** School of Life Sciences, University of Warwick, United Kingdom; Department of Statistics, University of Warwick, United Kingdom; Shanghai Institute of Materia Medica, Chinese Academy of Sciences, Shanghai, China; Big Data Institute, Nuffield Department of Population Health, University of Oxford, United Kingdom; Department for Continuing Education, University of Oxford, United Kingdom

**Keywords:** bacterial genomics, homologous recombination, Bayesian phylogenetics

## Abstract

Recombination is a major evolutionary force in bacteria, and ignoring it distorts phylogenies and can invalidate downstream analyses such as dating. We present ClonalFrame2, a new software for Bayesian phylogenetic inference from whole bacterial genomes that accounts for recombination. It replaces both ClonalFrame and ClonalFrameML while using the same evolutionary model, in which both mutation and recombination events occur on the branches of the clonal genealogy. The locations of recombination events are integrated out of the likelihood using a hidden Markov model, so the algorithm targets only the clonal genealogy and three recombination parameters. Ancestral sequences are recomputed whenever the tree changes and are used to propose efficient topology updates. We validated the method by analysing data-free datasets, which recovered the prior, and by comparing the approximated likelihood with the exact one on simulated data. Across 100 simulated datasets of 100 genomes each, the recombination parameters were accurately recovered, and trees inferred while accounting for recombination were many times more accurate than when ignoring it. In *Staphylococcus aureus* ST239, we showed that accounting for recombination gave a stronger temporal signal and a more robust dated phylogeny. In *Neisseria gonorrhoeae*, we identified interesting patterns of clustered recombination events happening repeatedly across the branches of the tree.

## INTRODUCTION

Recombination is an important evolutionary force in most bacterial species, following a process akin to gene-conversion in higher organisms, in which a relatively short fragment from a donor cell replaces the homologous fragment from a recipient cell (Vos and Didelot 2009; Didelot and Maiden 2010; Lin and Kussell 2017). Phylogenies are widely used to investigate bacterial population structures and epidemiology, but most standard phylogenetic tools ignore the effect of recombination which can distort the results (Schierup and Hein 2000; Hedge and Wilson 2014; Sakoparnig et al. 2021). Accounting for bacterial recombination when performing phylogenetic inference is therefore important to reconstruct an accurate tree, but also to study the recombination events themselves which often play a key evolutionary role, for example to adapt to new hosts and ecological niches (Sheppard et al. 2018), to escape vaccines (Golubchik et al. 2012; Croucher et al. 2013) or to acquire antibiotic resistance (Hanage et al. 2009; D’Aeth et al. 2021).

The currently most widely used tools to reconstruct phylogenies from whole bacterial genomes while accounting for bacterial recombination are Gubbins (Croucher et al. 2015) and ClonalFrameML (Didelot and Wilson 2015). Gubbins works by iterating between standard phylogenetic inference using raxml (Stamatakis 2014) and excluding genomic regions that are not well explained by this phylogeny. ClonalFrameML is a reimplementation of the original ClonalFrame software (Didelot and Falush 2007) which was designed to analyse multi-locus sequence typing (MLST) data (Maiden et al. 1998). ClonalFrame did not scale well to analysing whole genomes, whereas ClonalFrameML was developed specifically for this purpose, although this required compromises to be made compared to the original ClonalFrame. ClonalFrameML does not fully infer the phylogeny, but instead corrects the branch lengths of a tree given as input, while assuming that the topology is correct. The rationale for this approach was grounded in the observation that, in simulations, recombination mostly affects the branch lengths rather than the topology of a tree inferred without accounting for recombination (Hedge and Wilson 2014). Furthermore, ClonalFrameML uses maximisation of a target density, not Bayesian statistics like the original ClonalFrame, which is the most natural approach to capture uncertainties in the parameters.

ClonalFrame (Didelot and Falush 2007) and ClonalFrameML (Didelot and Wilson 2015) are based on the same evolutionary model in which the genomes are related through a tree known as the clonal genealogy. Mutation events occur on the branches of this tree as in any phylogenetic model, but in addition recombination events also occur at a different rate. Each time a recombination event happens, a short segment is affected in which each position is substituted with a fixed probability. This is a key difference with the ancestral recombination graph (ARG) model (Hudson 1983; Griffiths and Marjoram 1997) which tracks the ancestry of recombination donors, making inference very challenging. The ARG is the gold standard to model recombination within a population, for example in humans (Nielsen et al. 2025), viruses (Zhan et al. 2025) and indeed bacteria (Didelot et al. 2010; De Maio and Wilson 2017). Besides the advantage of computational simplicity, the ClonalFrame model also allows us to represent recombination events which may come from other populations than the one under study, for example if all the genomes under analysis belong to a single lineage which frequently imports material from other lineages or even other species (D’Aeth et al. 2021; Mourkas et al. 2022).

Here we present ClonalFrame2, a new software to perform Bayesian phylogenetic inference while accounting for recombination in whole bacterial genomes, replacing both ClonalFrame (Didelot and Falush 2007) and ClonalFrameML (Didelot and Wilson 2015). ClonalFrame2 borrows the best methodology from both its predecessors, including the same evolutionary model, as well as introducing its own methodology based on statistical and computational advances of the past few years. We demonstrate the use of ClonalFrame2 on a range of simulated and real genomic datasets.

## METHODS

### Model of evolution with bacterial recombination

The data is an alignment denoted *A* with *n* genomes and *L* sites. The clonal genealogy *T* is a parameter of the model, taking a value in the space of unrooted trees with 2n − 3 branches. Let *l*_*i*_ denote the length of branch indexed *i*. On the branches of the clonal genealogy, both mutation and recombination events can occur. Mutation events occur for each site at rate *l*_*i*_ whereas recombination events are initiated for each site at rate p*l*_*i*_. In other words, *p* is the rate of recombination relative to mutation (this parameter was previously denoted *R/θ* in Didelot and Wilson 2015). Let *δ* denote the mean of the Exponential distribution for the length of recombination events. Finally, the parameter *v* represents the evolutionary distance between donor and recipient during recombination.

The recombination model is a Hidden Markov Model (HMM) with two states: unimported (U) and imported (I). Since recombination events may overlap on a given branch of length *l*_*i*_ the expected length of a recombined region is longer than the length *δ* for a single event. A M/M/∞ queue can be used to describe the distribution of the lengths of recombined regions. The full distribution is complex (Roijers et al. 2007; Didelot et al. 2008), but its mean is *m =* (exp(p*l*_*i*_) − 1)/(p*l*_*i*_) and this distribution is approximated as Exponential(1/m) in order to retain the memoryless property. By combining this with the distribution of the length of unrecombined regions Exponential(p*l*_*i*_), the transition matrix of the HMM for branch *i* is obtained as:

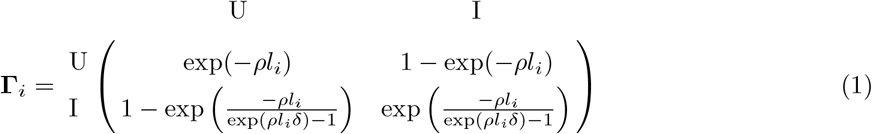

Each HMM is assumed to start at its steady state *π*_*i*_ obtained by solving ***π***_*i*_ *=* ***π***_*i*_ **Γ**, giving:

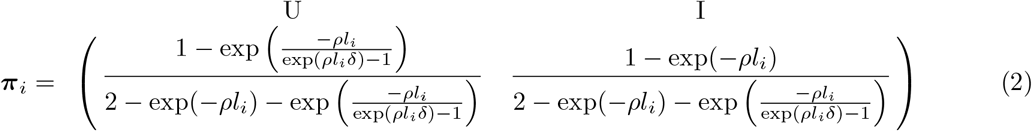

In the unimported state, the probability of genetic substitutions follows that of a Jukes-Cantor model (Jukes and Cantor 1969) of mutations with rate *l*_*i*_. In particular the probability of observing no

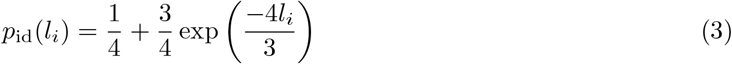

In the imported state the same mutation model is assumed but the rate is *v* instead of *l*_*i*_. For example, the probability of observing no difference is given by *p*_id_(*v*) using Equation 3.

### Likelihood computation

The likelihood is computed by considering only a single set of ancestral sequences for a given clonal genealogy, corresponding to the maximum likelihood reconstruction of joint sequences (Pupko et al. 2000). This represents an approximation compared to the likelihood that would be obtained by integrating over the space of possible ancestral sequences, the effect of which will be investigated. The ancestral state reconstruction is not constant but instead recomputed every time the tree changes as will be described later.

Since for a given tree the ancestral sequences are known exactly, the overall likelihood is the product of the likelihood of each branch:

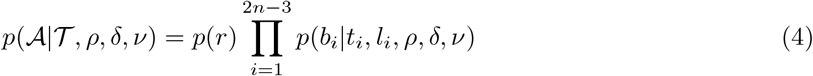

where *b*_*i*_ and *t*_*i*_ are the sequences at the bottom and the top of the branch indexed *i*, respectively. *p*(*r*) is the probability of the sequence at the root of the tree, for example under the Jukes-Cantor model this is simply *p*(*r*) = 1/4^*L*^.

To calculate the likelihood of a branch *p*(*b*_*i*_|*t*_*i*_, *l*_*i*_, *ρ, δ, ν*), the HMM forward algorithm is used (Rabiner 1998). Durbin et al. 1989; Note that this calculates the likelihood integrated over the recombination events. This way sampling over the space of recombination events is avoided compared to previous implementations (Didelot and Falush 2007; Didelot and Wilson 2015). It is still possible to output recombination events including their uncertainty using the forward-backward sampling algorithm (Rabiner 1989; Durbin et al. 1998), but this is done a-posteriori and separately from the main inference procedure.

The calculation of the integrated likelihood is as follows:

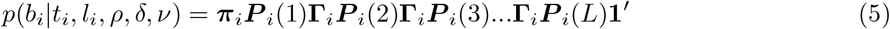

where **Γ**_*i*_ and ***π***_*i*_ are defined in Equations 1 and 2, respectively. The matrix ***P*** _*i*_(*j*) is a 2x2 diagonal matrix containing the probability of observing the data at site *j* for branch *i* in the unimported and imported states, respectively. In particular, when the same nucleotide is observed at the top and bottom of a branch, this is equal to:

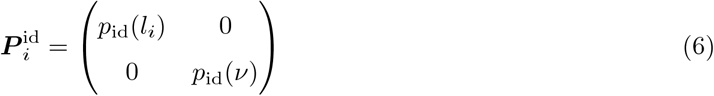

where *p*_id_(*x*) is defined in Equation 3. Since typically many sites have identical sequences at the top and bottom of a branch, the computation in Equation 5 can be optimised by pre-computing 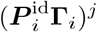 for every value of *j* from 1 to 100 for example. Each value is obtained from the previous one simply by multiplying with 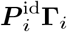. Note that this is an exact optimisation, not an approximation, and does not assume in particular that only a single HMM transition happened between polymorphic sites.

The likelihood computation is the most computationally expensive part of the inference algorithm. The forward algorithm (Equation 5) has complexity *O*(*L*) and needs to be performed for each branch so that the overall complexity of the likelihood computation is *O*(*Ln*). However, since the likelihood is decomposed across branches (Equation 4) this calculation is parallelised over branches using OpenMP (Dagum and Menon 1998).

### Bayesian inference

As explained above, the ancestral sequences are fixed given the tree and the recombination events are integrated out from the likelihood. Therefore the Bayesian inference targets only the clonal genealogy, and the three parameters of the recombination process, making inference very effcient compared to previous implementations which had to mix or optimise over the very large parameter space of import locations (Didelot and Falush 2007; Didelot and Wilson 2015). The posterior target is decomposed using Bayes rule and the conditional independence of the model parameters as follows:

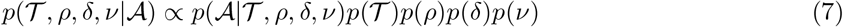

The prior on the clonal genealogy is that all of the (2*n −* 5)!! topologies are equally likely and the branch lengths follow a Gamma(*α, θ*) prior, so that:

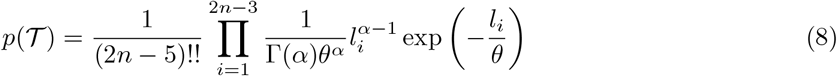

The values of the hyper-parameters *α* and *θ* can be selected by the user. Likewise, the priors on *ρ, δ* and *ν* are Gamma distributions with parameters that can be selected by the user. The default values are *α* = 1 and *θ* = 10^*−*4^ for the branch lengths, shape parameters for *ρ, δ* and *ν* all equal to 1 and scale parameters of 1, 500 and 0.05, respectively.

Inference is done using a Markov Chain Monte-Carlo (MCMC) algorithm (Gilks et al.1996). The update of the three parameters of the recombination process *ρ, δ* and *ν* uses an adaptive Metropolis-Hastings update with a Normal proposal kernel and acceptance target 0.234 (Andrieu and Thoms 2008; Spencer 2021). The branch lengths of the clonal genealogy are also updated using the same strategy. Updates of the tree topology are proposed using a version of subtree pruning and regrafting (SPR) guided by the ancestral sequences, similar to (Wilson and Balding 1998). This is a big advantage of having ancestral sequences since otherwise unguided SPR would have low chance of acceptance (Höhna and Drummond 2012; Zhang et al. 2020). The guiding process uses the ancestral sequences before the tree move but the likelihood of the newly proposed tree is calculated using newly generated ancestral sequences. This SPR is therefore inspired by a Gibbs move (Gelfand and Smith 1990), in the sense that it would almost be targeting the correct conditional distribution and have an acceptance probability of one if the ancestral sequences were unchanged. Since they are changed the move is a Metropolis-Hastings, and not a Gibbs move, but it should still have a high acceptance probability. The proposed pruning node *i* is selected uniformly, and the proposed regrafting sister node *j* is selected amongst the remaining nodes with probability that accounts for the number of differences *d*(*i, j*) between them as follows:

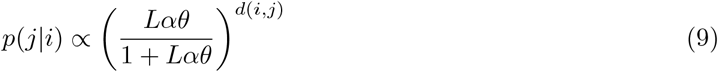

This geometric probability corresponds to the integrated probability for a branch of length *l* sampled from an Exponential distribution with mean *αθ* (ie with the same mean as a Gamma(*α, θ*)) of having its number of substitutions distributed as Poisson(*Ll*) equal to *d*(*i, j*). In particular, the value of *p*(*j*|*i*) decreases as *d*(*i, j*) increases, so that this proposal mechanism tends to propose regrafting sister nodes that have similar ancestral sequences to the pruning node. Once the nodes *i* and *j* are selected, a new node *n* is created with children *i* and *j* and mother *f* equal to the previous mother of *j*. The three new branch lengths *l*(*n, i*), *l*(*n, j*) and *l*(*n, f*) between *n* and (*i, j, f*) are proposed by first computing the differences *d*(*i, f*), *d*(*i, j*) and *d*(*j, f*). Figure S1 illustrates that these distances are expected to have the following additive properties as follows:

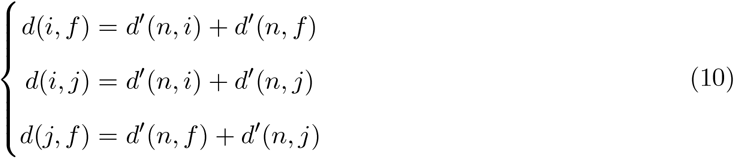

Solving these three equations gives the expected number of differences on each of the three new branches:

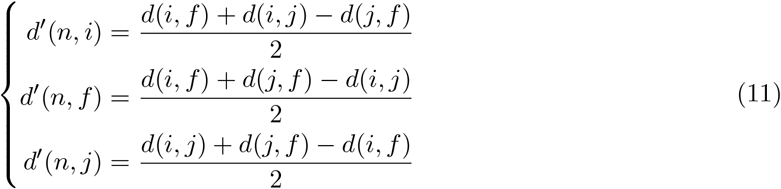

Likewise, the newly created branch where pruning happened which now connects the previous mother *p* of *i* and its previous sister *b* has expected number of differences *d*^*’*^(*p, b*) = *d*(*p, b*). Each of these four branches has a length with prior Gamma(*α, θ*) and a number *d*^*’*^ of differences are expected out of *L* sites, of which a proportion 1/(1 + *ρδν*) are expected to be caused by mutation rather than recombination. We therefore use the Gamma-Poisson conjugacy to propose the new branch length *l*^*’*^ as follows:

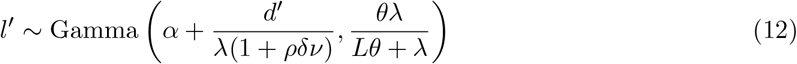

Note that setting *λ* = 1 would be using the exact conjugacy but since the move is not exactly Gibbs, we use a higher value to make the proposal distribution have heavier tails, typically *λ* = 2 but this value can be fine-tuned to obtain a good acceptance probability. The choice of this value of *λ* and of the exact form of the proposal distributions in Equations 9 and 12 does not affect the target of the MCMC, only its convergence and mixing properties.

### Implementation

All methods described in this article are available at https://github.com/xavierdidelot/ClonalFrame2. The implementation is written in C++, and OpenMP is used for the parallelisation (Dagum and Menon 1998). By default the clonal genealogy is initiated using the Neighbor-Joining algorithm (Saitou and Nei 1987) . Each time an updated tree is proposed, the ancestral states are computed using the maximum likelihood algorithm of Pupko et al. (2000) before its likelihood is computed using Equation 4. Any gap in the alignment is treated as missing data and integrated out in the ancestral state reconstruction. An accompanying R package entitled RClonalFrame2 provides an R interface to run ClonalFrame2 and to analyse results. This relies on the phylogenetic tools from packages ape (Paradis and Schliep 2019) and phangorn (Schliep 2011). All results were computed on a standard laptop from 2018 with processor 2.9 GHz 6-Core Intel Core i9 and memory 16 GB 2400 MHz DDR4.

## RESULTS

### Summary and validation of the analytical approach

Our evolutionary model with bacterial recombination (Didelot and Falush 2007; Didelot and Wilson 2015) is illustrated in Figure 1, showing all events that occurred on all branches of a simulated clonal genealogy. There are several points worth noticing that are direct consequences of the model. First, there are more mutation and recombination events on longer branches. Second, in recombined regions there are more substitutions than in unrecombined regions, because here a value of *ν* = 0.01 was used which is ten times more than the average branch length of 0.001. Third, longer branches tend to have longer recombined regions, due to the higher probability of several recombination events overlapping.

**Figure 1.**
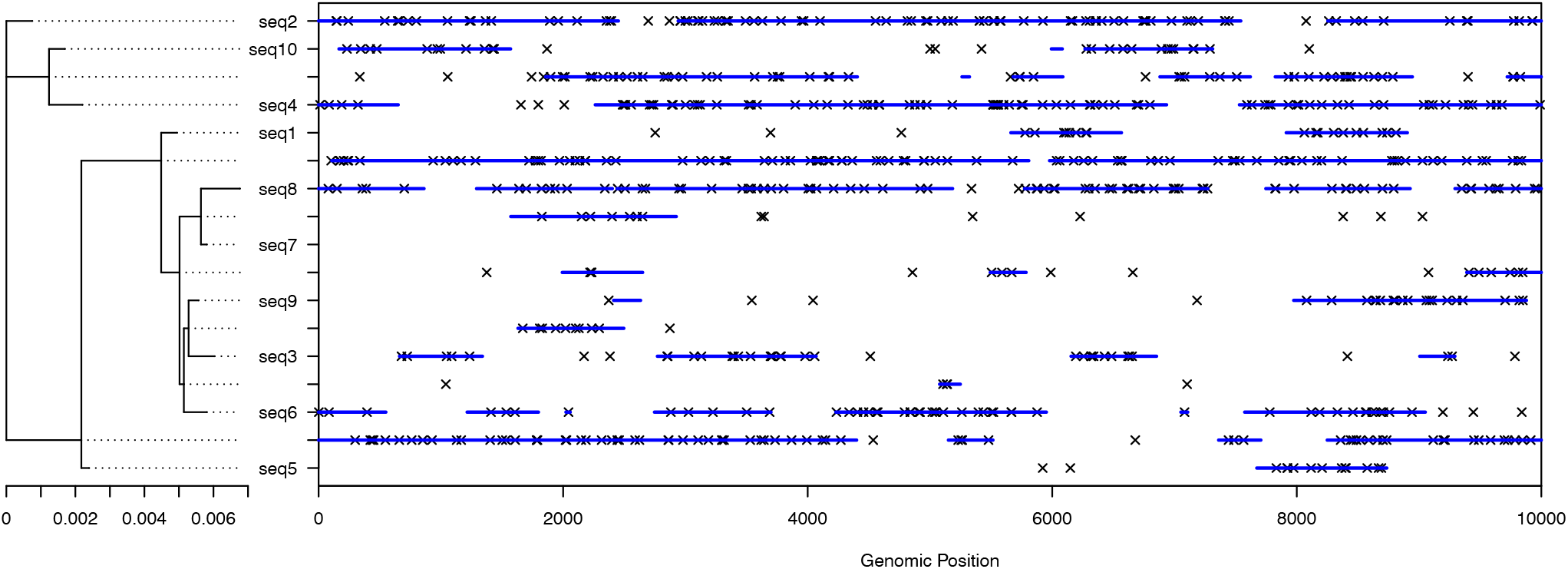
Illustration of the evolutionary model with bacterial recombination. The clonal genealogy is shown on the lefthand side. For each branch of the clonal genealogy, there is a horizontally aligned row on the righthand side, showing the evolutionary events that occurred on that branch. The recombination events are shown as blue bars, while substitutions are shown as black crosses.

Bayesian inference under this model is performed exactly with one exception: when computing the likelihood of a given tree, the ancestral sequences are conditioned to be equal to their joint maximum likelihood estimate (Pupko et al. 2000). This approach is similar to the one used in ClonalFrameML (Didelot and Wilson 2015) where ancestral sequences were computed for a single input tree and considered fixed throughout the algorithm. Here however the ancestral sequences are recomputed every time the tree is changed, which makes it more accurate. To show that the approximation is relatively mild, we simulated a tree with *n* = 20 leaves from the prior (Figure 2A) and from this we simulated genomes of length *L* = 10^6^ without recombination. In this condition, we can compute the full likelihood using Felsenstein pruning (Felsenstein 1981) and compare it with our approximate conditioned likelihood. We performed this comparison for 100 trees (two examples are shown in Figure 2B and C), each of which was a slightly randomised version of the correct tree (Figure 2A). Figure 2D shows that there is little difference between the full likelihood and our conditional likelihood. Further evidence that this approximation does not invalidate results will come from application to simulated datasets.

**Figure 2.**
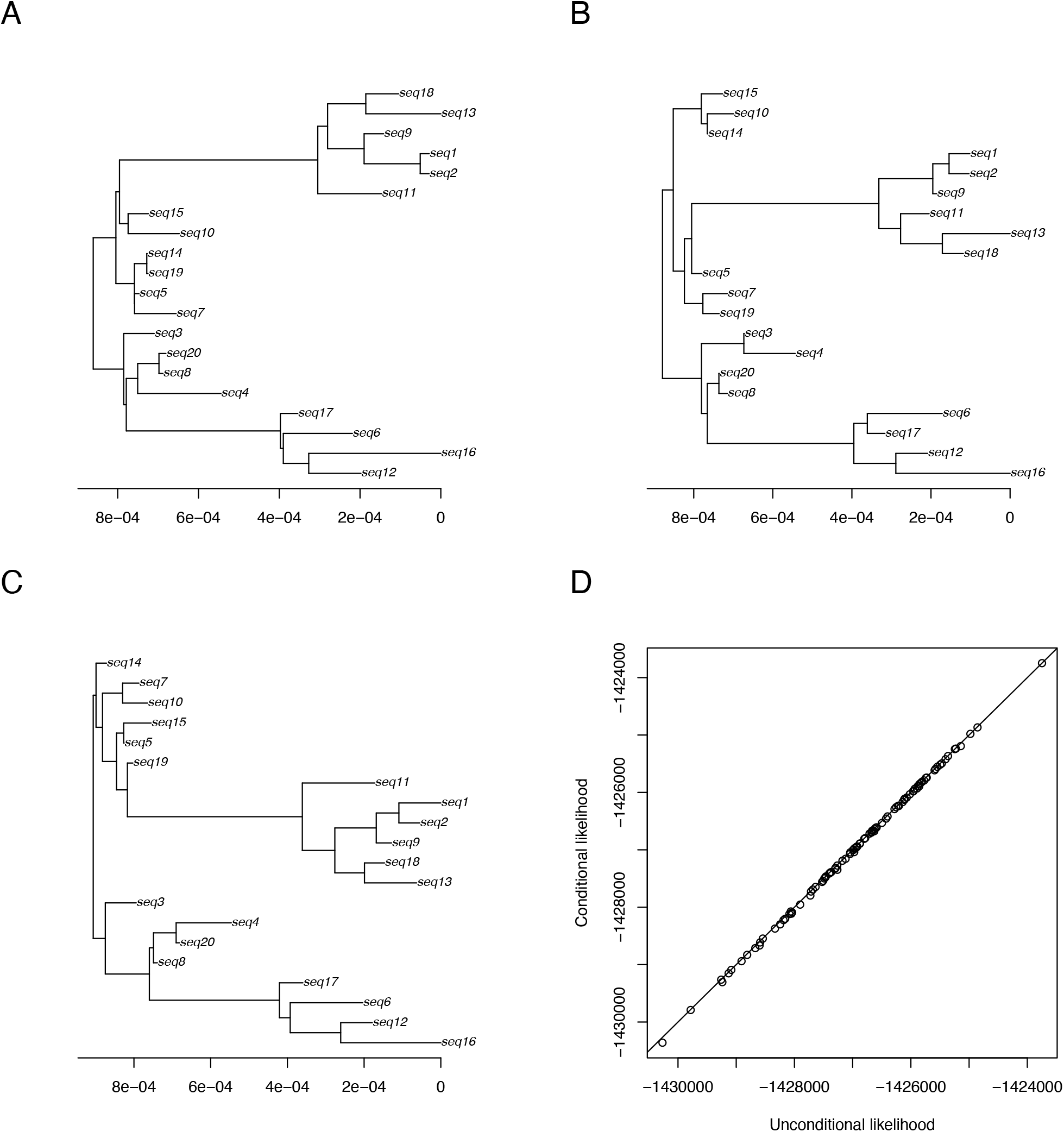
Comparison of conditional and full likelihoods. (A) Simulated phylogeny with twenty leaves. (B) An example of randomised tree. (C) Another example of randomised tree. (D) Comparison of the conditional and full likelihoods for 100 randomised trees.

To ensure that the MCMC is working correctly, inference was performed on a dataset with *n* = 100 sequences of length *L* = 0. In this case, since there is no sequence data, the MCMC is expected to explore the priors for the branch lengths and for the parameters *ρ, δ* and *ν* which were Gamma-distributed with means 10^*−*3^, 1, 500 and 0.05 respectively, and standard deviations equal to half of the means. Note that this is not the default prior (which has shape *α* = 1 and therefore means equal to their standard deviations) to avoid the special case in which the Gamma reduces to an Exponential distribution and therefore provide a more general test. We found that this is indeed the case for the parameters *ρ*,δand *v* (Figure 3A, B and C), for the sum of the branch lengths of the tree (Figure 3D), for a single branch length (Figure 3E) and for the value of the tree prior which is distributed as expected in Equation 8 (Figure 3F). A majority-rule consensus tree was computed and found to be entirely star-like as expected (Figure S2). We also performed smaller runs with *n* = 5 and *n* = 6 for which a chi-square test on the one-dimensional contingency table of sampled tree topologies was used to check that the tree topologies are sampled uniformly. This test of the MCMC when using no sequence data validates all parts of the MCMC except for the likelihood function (Equation 4). To ensure that it is correct, we checked that our C++ implementation returns exactly the same values as a separate implementation in R based on the ancestral state reconstruction performed by the *phangorn* package (Schliep 2011). We also tested both in R and in C++ that the HMM optimisation described after Equation 6 was correctly implemented by making sure that the values of the likelihoods were the same with and without this optimisation.

**Figure 3.**
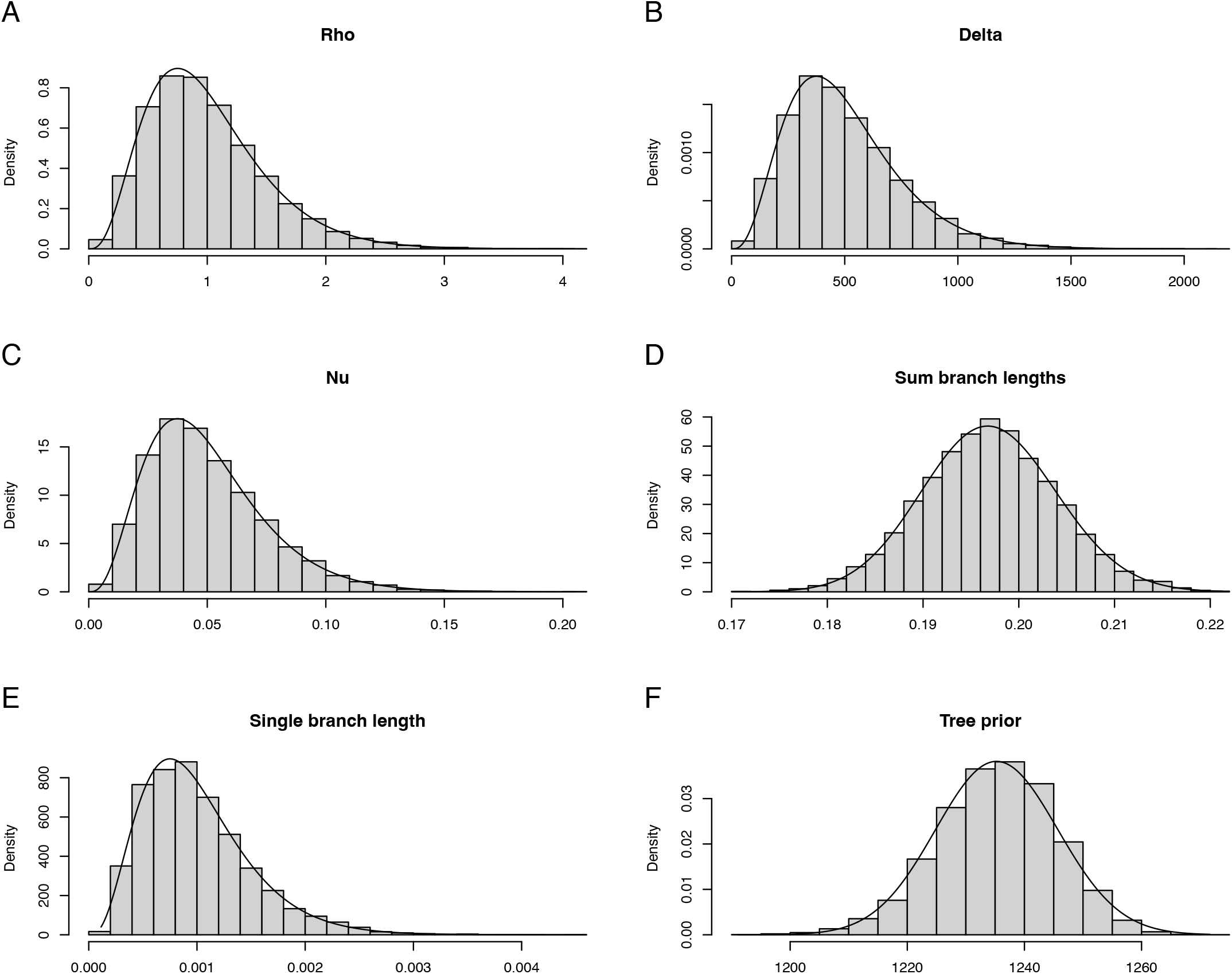
Validation of the MCMC by analysing without any data and checking that the posterior samples (histograms) follow their prior distributions (lines).

### Application to simulated datasets

We simulated 100 separate datasets under our model, each of which was made of *n* = 100 genomes of length *L* = 10^6^. The values of the parameters were chosen at random uniformly on a log_10_ scale in the following ranges: *ρ* ∈ [0.01, 1],δ∈ [100, 1000] and *v* ∈ [0.01, 0.1]. For each simulation, we performed Bayesian inference by running our MCMC algorithm for a total of 10^5^ iterations. The starting point of each run was *ρ* = 0.1,δ= 500, *v* = 0.05 and the tree was initiated using the Neighbor-Joining algorithm (Saitou and Nei 1987). Each run took around 30 minutes on a standard laptop computer, with runs for larger values of the relative impact of recombination r/m=*ρdv* taking longer due to having more polymorphisms caused by recombination. The parameters inferred in each run are compared with the parameters used in the simulation in Figure 4. The mean inferred value for each parameter is always close to the correct value, and the 95% credible interval almost always contains the correct value. The coverage frequencies are 93/100 for *ρ*, 94/100 forδand 93/100 for *v*, which is approximately as expected for the 95% credible intervals.

**Figure 4.**
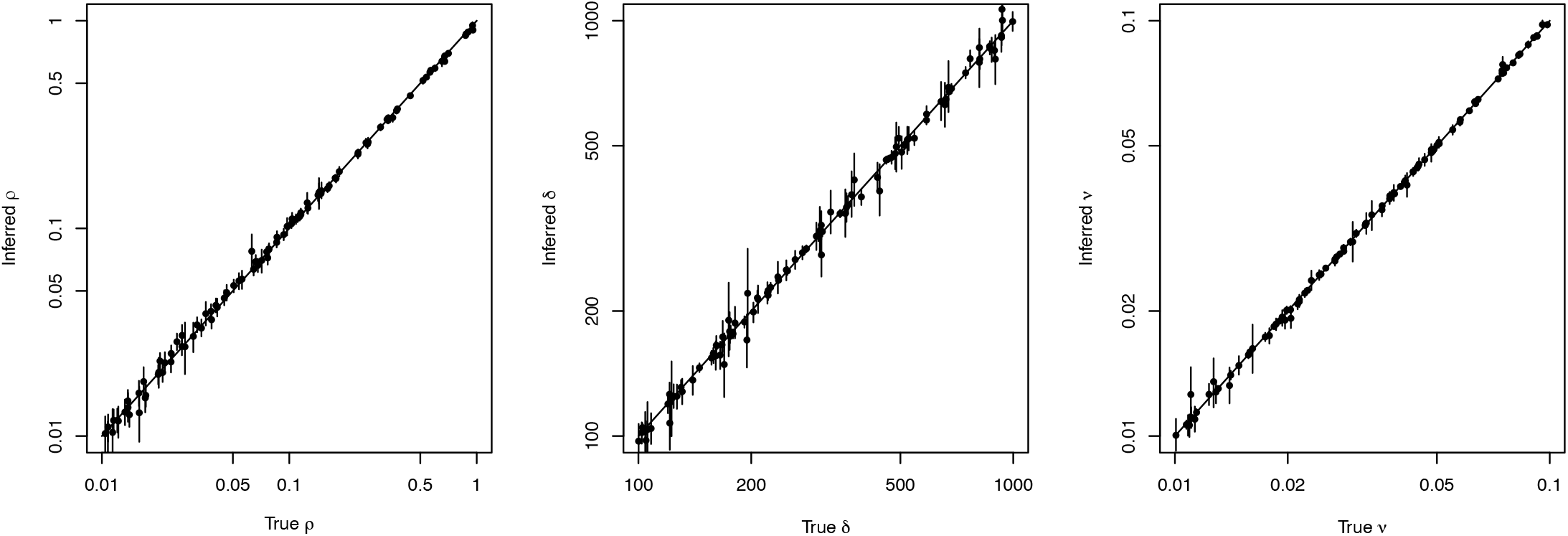
Comparison of the correct (x-axis) and inferred (y-axis) values of the three parameters *ρ, δ* and *v* for 100 simulated datasets. The bars correspond to the 95% credible intervals.

In order to confirm that accounting for recombination improves the phylogenetic inference of the clonal genealogy, we performed a separate run for each dataset in the same conditions as above but with the recombination rate forced equal to *ρ* = 0. We then computed the branch score distance (Kuhner and Felsenstein 1994) between sampled inferred trees and the correct tree. When recombination was inferred, this distance was on average 1.92 × 10^−4^, which is about the same as the average distance of 1.90 × 10^−4^ between two sampled trees. In other words, the correct tree is about as distant from a sampled tree as any two sampled trees are from each other, as would be expected when Bayesian inference is consistent with calibration (Cook et al. 2006; Talts et al. 2020). On the other hand, when recombination was ignored, the average distance between a sampled tree and the correct tree was 5.39 × 10^−3^ which is about 28 times higher than when recombination was inferred. This is also about 16 times more than the average distance of 3.33 × 10^−4^ between two sampled trees, which strongly suggests that the correct tree is not in the posterior distribution when recombination is ignored. The average Robinson-Foulds distance (Robinson and Foulds 1981) between the true tree and trees sampled ignoring recombination was 1.58 compared to 1.42 when recombination was accounted for. Since this distance only depends on the topology of the trees, it shows that the topological inference is slightly improved when accounting for recombination, but that there is a more important improvement in the branch lengths, as previously suggested (Hedge and Wilson 2014). The overall inference of the phylogeny is therefore more accurate when recombination is correctly modelled and accounted for.

### Application to Staphylococcus aureus ST239

We reanalysed the dataset of *Staphylococcus aureus* ST239 originally published by Harris et al. (2010), in which the evolutionary history and global dissemination of a multidrug resistant hospital-associated pathogen was reconstructed using whole genome sequencing. We obtained an alignment of 59 genomes isolated between 1982 and 2007 by applying Snippy version 4.6.0 (https://github.com/tseemann/snippy) to each genome against the complete reference genome TW20, which is a member of ST239 (Holden et al. 2010). We ran ClonalFrame2 on this dataset for 10^6^ iterations, discarding the first half as burn-in and sampling the remainder every 500 iterations to obtain posterior samples of size 1000. Each run took approximately an hour on a standard laptop. Two settings were applied: the default setting accounting for recombination and an alternative setting ignoring recombination, obtained by forcing the recombination rate to *ρ* = 0. In each setting, the Gelman-Rubin diagnostic (Gelman and Rubin 1992; Brooks and Gelman 1998) was lower than 1.1 and the effective sample size statistic implemented in the R package coda (Plummer et al. 2006) was greater than 200 for the inferred continuous parameters *ρ, δ, v* and the sum of branch lengths in the tree. The approximate effective sample size of the tree topology was also greater than 200 as computed by the R package rwty (Warren et al. 2017). In the default setting, the parameters were inferred to be *ρ* = 0.11 [0.09; 0.12], *δ* = 261 [235; 290] and *v* = 0.036 [0.035; 0.038] so that the relative effect of recombination (versus mutation) was equal to r/m=*ρ δv* = 1.00 [0.88; 1.13]. All values in square brackets correspond to the 95% credible intervals. The effect of recombination is higher than usually found in *S. aureus* as a whole, but in good agreement with previous estimates for the ST239 lineage (Castillo-Ramírez et al. 2012; Everitt et al. 2014; Didelot and Wilson 2015).

In each setting (with and without recombination), the medoid tree was extracted corresponding to the sampled tree with the average smallest branch score distance (Kuhner and Felsenstein 1994) to all other sampled trees. The tree without recombination had a weaker temporal signal compared to the tree with recombination, as assessed using the correlation of root-to-tip distances with sampling dates *(R*^*2*^ = 0.45 vs *R*^*2*^ = 0.82, Figure 5). Both trees were dated using BactDating (Didelot et al. 2018) under the additive relaxed clock model (Didelot et al. 2021). In the analysis ignoring recombination, the clock rate was estimated to be *μ* = 18.8 [13.6; 25.0] substitutions per year, the relaxation parameter *σ* = 31.8 [23.0;42.9] and the date of the root 1954 [1938; 1966]. In the analysis accounting for recombination, the clock rate was estimated to be *μ* = 9.1 [7.2; 11.4] substitutions per year, the relaxation parameter *σ* = 3.7 [2.2; 6.0] and the date of the root 1962 [1952; 1969]. The higher value of the relaxation parameter for the tree ignoring recombination is a telling sign that this analysis worked less well, and this resulted in a wider range for the dating of the root and other nodes (Figure 5). We applied DiagnoDating (Didelot et al. 2026) to assess the validity of these dating analyses and found that when recombination was ignored the posterior predictive check raised an issue *(p* = 0.008) whereas when recombination was accounted for this was not the case (p = 0.373). In conclusion, these results demonstrate that the tree accounting for recombination is more accurate than the tree ignoring recombination. This application also showcases the importance of correcting a tree for recombination before performing a dating analysis and any further analysis that depends on the dated phylogeny (Didelot and Parkhill 2022).

**Figure 5.**
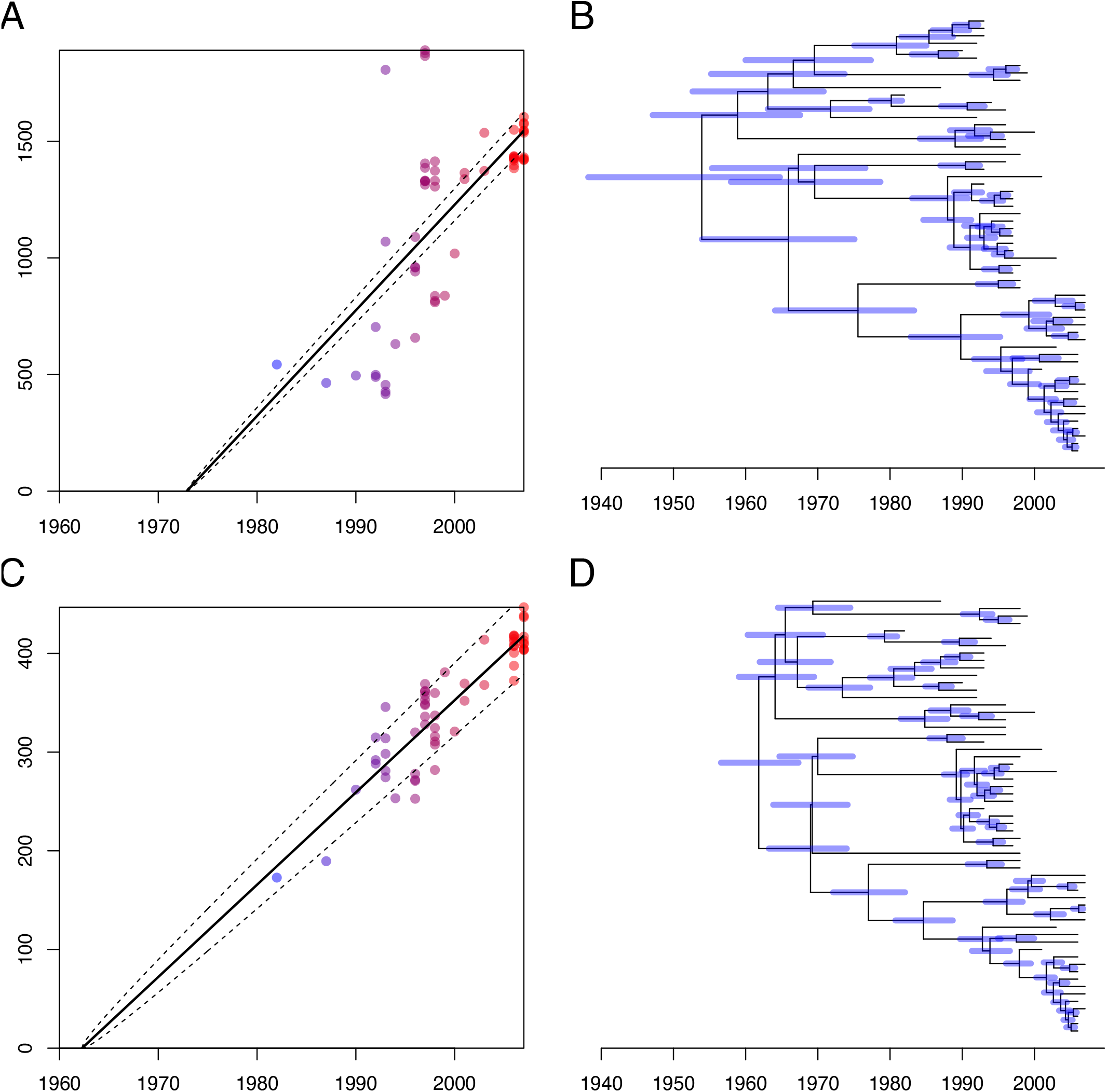
(A) Root-to-tip distance analysis for the tree ignoring recombination. (B) Dating of the tree ignoring recombination. (C) Root-to-tip distance analysis for the tree accounting for recombination. (D) Dating of the tree accounting for recombination.

### Application to *Neisseria gonorrhoeae* cefixime resistant clade 1

The genomic epidemiology of *Neisseria gonorrhoeae* was previously studied by Grad et al. (2014) which described that two separate clades resistant to cefixime had emerged in the late 1990s and subsequently spread throughout the USA. A total of 104 genomes were found to belong to the cefixime resistant cluster 1, and we generated an alignment of these genomes using Snippy against the complete sequence of the reference genome TCDC-NG08107 (Chen et al. 2011). Running ClonalFrame2 on this dataset for 10^6^ iterations took approximately 3 hours on a standard laptop. Good convergence and mixing properties were confirmed as described in the previous example. The parameter estimates were *ρ* = 0.42 [0.37; 0.48], *δ* = 206 [184;231] and *v* = 0.035 [0.032;0.038], so that the relative effect of recombination versus mutation is r/m=*ρ δv* = 3.08 [2.69; 3.55].

Figure 6 shows the medoid tree from the posterior sample, as well as the posterior probability of recombination for each branch of the tree and each position along the genome. Overall the distribution of recombination events across branches and genomic locations looks fairly sparse and uniform, but there are a few striking exceptions. Some locations are clearly marked as recombinant over more branches than would be expected just by chance. The most recombinant location was position 671,500bp with a sum of per-branch recombination probabilities equal to 11.44, coinciding with the gene encoding protein ADV07653 in the annotation of TCDC-NG08107 (Chen et al. 2011). This is annotated as a membrane protein in the original annotation and as a DUF456 domain-containing protein in other genomes. There was also a wider cluster of recombination events around position 1.1Mbp where several genes encoding phage associated proteins have been reported (Chen et al. 2011). Another smaller cluster was found between positions 1.27Mbp and 1.33Mbp (Figure 6), which encodes the TbpA and TbpB proteins responsible for iron transport from human transferrin and have been previously described as a recombination hotspot in *N. gonorrhoeae* and other bacterial pathogens (Harrison et al. 2008; Yahara et al. 2016). These two regions were not inferred to be continuously recombinant, but included positions with sums of recombination probabilities up to 7.99 and 9.51, respectively. The position 1.958Mbp had a sum of recombination probabilities equal to 5.14 and corresponded to the annotated gene ADV08938 which encodes the outer membrane porin PorB which is a well known hotspot of recombination in *N. gonorrhoeae* (Yahara et al. 2016; Manoharan-Basil et al. 2023).

**Figure 6.**
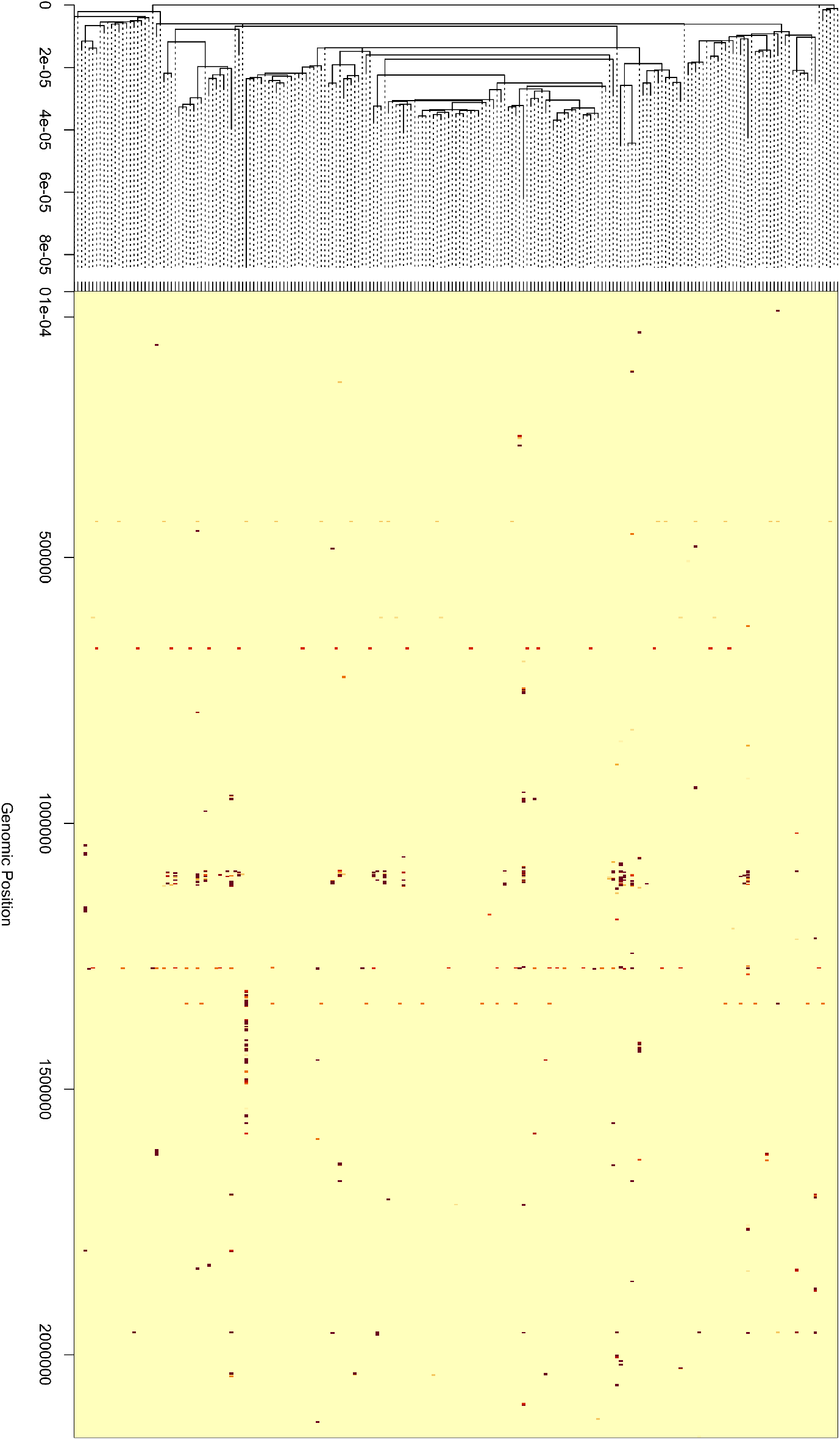
Phylogenetic tree and recombination events inferred on each branch in the *Neisseria gonorrhoeae* analysis. Darker red indicates a higher posterior probability of recombination for a given branch and genomic location.

Finally, it is remarkable that one genome denoted GCGS143 has a long cluster of recombination between positions 1.3Mbp and 1.5Mbp, resulting in the longest terminal branch in the inferred clonal genealogy (Figure 6). This cluster is about 200kbp in length but broken by several clonal regions, and there are several similar but smaller broken clusters of recombination found in other genomes, especially in the previously mentioned highly recombinant region around position 1.1Mbp. The main route of recombination in *N. gonorrhoeae* is natural transformation, followed by RecA-mediated homologous recombination of imported DNA into the chromosome (Hamilton and Dillard 2006). This is also true of *Helicobacter pylori* for example, in which very similar patterns of long recombination events interspersed with unrecombined sequences have been previously reported (Kennemann et al. 2011; Krebes et al. 2014; Bubendorfer et al. 2016).

## DISCUSSION

We have presented ClonalFrame2, a Bayesian method for phylogenetic inference from whole bacterial genomes that accounts for recombination. It retains the evolutionary model of ClonalFrame (Didelot and Falush 2007) and ClonalFrameML (Didelot and Wilson 2015) illustrated in Figure 1, but its inference differs in several ways. Recombination events are integrated out of the likelihood using the HMM forward algorithm, so the MCMC targets only the clonal genealogy and three recombination parameters. A similar difference is described for example between the original version of a spatiotemporal epidemic model (Knorr-Held and Richardson 2003) and its recent reimplementation (Adeoye et al. 2026). Ancestral sequences are reconstructed by maximum likelihood and recomputed whenever the tree changes, unlike ClonalFrameML which assumed a fixed tree and ancestral sequences. Topology updates use an SPR move guided by the ancestral sequences. In simulations, the posterior distributions of the recombination parameters were close to the true values (Figure 4), and ignoring recombination gave trees about 28 times further from the truth than those accounting for it. In the reanalysis of *S. aureus* ST239 (Harris et al. 2010), being aware of recombination gave a stronger temporal signal and a more robust dating analysis (Figure 5). In the reanalysis of *N. gonorrhoeae* (Grad et al. 2014), the posterior recombination probabilities highlighted loci that recombine repeatedly across the tree (Figure 6) and that corresponded to previously described recombination hotspots.

Our runs took between 1 and 3 hours on a standard laptop for datasets around a hundred genomes. We expect ClonalFrame2 to be practical for datasets of up to a few thousand genomes. The computing time scales approximately linearly per MCMC iteration (Equation 4), but larger numbers of genomes may require more MCMC iterations (Whidden and Matsen 2015). For larger collections, we recommend splitting the dataset into clades and analysing each separately. This is not wasteful, since the long branches separating clades are rarely of interest. Each clade contains fewer genomes and also fewer polymorphisms, which makes the computation much more efficient. Working with less diverse clades also makes the model more accurate, since it is fundamentally a model of imports from external sources. The origin of the imported material and the flux of recombination between clades can then be investigated as a post-processing step (Didelot et al. 2011; Sheppard et al. 2013; Ozer et al. 2019).

Like its predecessors, ClonalFrame2 models recombination as the import of short segments from outside the clonal genealogy, and it does not track the ancestry of donors as in an ARG (Hudson 1983; Griffiths and Marjoram 1997). This keeps inference tractable and allows imports from other populations or species, but it is also a misspecification when most recombination occurs between members of the sampled population. Our simulations were generated under the inference model, so they do not test this case. We would expect within-population recombination to produce imported segments that are less divergent than the model assumes, which would make them harder to detect and more likely to be absorbed into branch lengths or topology. We previously showed this to be the case when simulating under an ARG model and inferred with ClonalFrameML (Didelot and Wilson 2015). As a first step towards resolving this issue, the recombination model could be extended beyond the two states clonal vs recombinant. A third state would allow two classes of imported segment, with lower and higher values of *v*, possibly capturing recombination from within and from outside the population of interest. The design of such model extensions would best be guided by Bayesian model comparison, for instance by computing a Bayes factor between a model with two states and one with three. Marginal likelihoods for phylogenetic models can be estimated by thermodynamic integration (Lartillot and Philippe 2006) or stepping-stone sampling (Xie et al. 2011), whereas reversible-jump MCMC (Hastie and Green 2012) is an alternative that avoids estimating marginal likelihoods altogether.

## References

Adeoye M, Didelot X, Spencer SE. 2026. Bayesian spatio-temporal modelling for infectious disease outbreak detection. Epidemics. 54:100879.

Andrieu C, Thoms J. 2008. A tutorial on adaptive MCMC. Statistics and Computing. 18:343–373.

Brooks SPB, Gelman AG. 1998. General methods for monitoring convergence of iterative simulations. Journal of computational and graphical statistics. 7:434–455.

Bubendorfer S, Krebes J, Yang I, Hage E, Schulz TF, Bahlawane C, Didelot X, Suerbaum S. 2016. Genome-wide analysis of chromosomal import patterns after natural transformation of Helicobacter pylori. Nature Communications. 7:11995.

Castillo-Ramírez S, Corander J, Marttinen P, Aldeljawi M, Hanage WP, Westh H, Boye K, Gulay Z, Bentley SD, Parkhill J, et al. (12 co-authors). 2012. Phylogeographic variation in recombination rates within a global clone of methicillin-resistant Staphylococcus aureus. Genome biology. 13:R126.

Chen CC, Hsia KC, Huang CT, Wong WW, Yen MY, Li LH, Lin KY, Chen KW, Li SY. 2011. Draft Genome Sequence of a Dominant, Multidrug-Resistant Neisseria gonorrhoeae Strain, TCDC-NG08107, from a Sexual Group at High Risk of Acquiring Human Immunodeficiency Virus Infection and Syphilis. Journal of Bacteriology. 193:1788–1789.

Cook SR, Gelman A, Rubin DB. 2006. Validation of Software for Bayesian Models Using Posterior Quantiles. Journal of Computational and Graphical Statistics. 15:675–692.

Croucher NJ, Finkelstein JA, Pelton SI, Mitchell PK, Lee GM, Parkhill J, Bentley SD, Hanage WP, Lipsitch M. 2013. Population genomics of post-vaccine changes in pneumococcal epidemiology. Nature genetics. 45:656–663.

Croucher NJ, Page AJ, Connor TR, Delaney AJ, Keane JA, Bentley SD, Parkhill J, Harris SR. 2015. Rapid phylogenetic analysis of large samples of recombinant bacterial whole genome sequences using Gubbins. Nucleic Acids Research. 43:e15.

D’Aeth JC, van der Linden MP, McGee L, de Lencastre H, Turner P, Song JH, Lo SW, Gladstone RA, Sá-Leão R, Ko KS, et al. (15 co-authors). 2021. The role of interspecies recombination in the evolution of antibiotic-resistant pneumococci. eLife. 10:1–35.

Dagum L, Menon R. 1998. OpenMP: An industry standard API for shared-memory programming. IEEE Computational Science and Engineering. 5:46–55.

De Maio N, Wilson DJ. 2017. The Bacterial Sequential Markov Coalescent. Genetics. 206:333–343.

Didelot X, Bowden R, Street T, Golubchik T, Spencer C, McVean G, Sangal V, Anjum MF, Achtman M, Falush D, et al. (11 co-authors). 2011. Recombination and Population Structure in Salmonella enterica. PLoS Genetics. 7:e1002191.

Didelot X, Carson J, Ribeca P, Volz E. 2026. DiagnoDating: Diagnostics for dated phylogenies in microbial population. Molecular Biology and Evolution. 43:msag093.

Didelot X, Croucher NJ, Bentley SD, Harris SR, Wilson DJ. 2018. Bayesian inference of ancestral dates on bacterial phylogenetic trees. Nucleic Acids Research. 46:e134–e134.

Didelot X, Falush D. 2007. Inference of bacterial microevolution using multilocus sequence data. Genetics. 175:1251–66.

Didelot X, Lawson DJ, Darling AE, Falush D. 2010. Inference of homologous recombination in bacteria using whole-genome sequences. Genetics. 186:1435–49.

Didelot X, Maiden MCJ. 2010. Impact of recombination on bacterial evolution. Trends in Microbiology. 18:315–322.

Didelot X, Parkhill J. 2022. A scalable analytical approach from bacterial genomes to epidemiology. Philosophical Transactions of the Royal Society B: Biological Sciences. 377:20210246.

Didelot X, Siveroni I, Volz EM. 2021. Additive uncorrelated relaxed clock models for the dating of genomic epidemiology phylogenies. Molecular Biology and Evolution. 38:307–317.

Didelot X, Taylor JE, Watkins JC. 2008. A Duality Identity between a Model of Bacterial Recombination and the Wright-Fisher Diffusion. Markov Processes and Related Topics. 4:315–324.

Didelot X, Wilson DJ. 2015. ClonalFrameML: Efficient Inference of Recombination in Whole Bacterial Genomes. PLoS Computational Biology. 11:e1004041.

Durbin R, Eddy SR, Krogh A, Mitchison G. 1998. Biological Sequence Analysis: Probabilistic Models of Proteins and Nucleic Acids. Cambridge: Cambridge University Press.

Everitt RG, Didelot X, Batty EM, Miller RR, Knox K, Young BC, Bowden R, Auton A, Votintseva A, Larner-Svensson H, et al. (19 co-authors). 2014. Mobile elements drive recombination hotspots in the core genome of Staphylococcus aureus. Nature Communications. 5:3956.

Felsenstein J. 1981. Evolutionary trees from DNA sequences: A maximum likelihood approach. Journal of Molecular Evolution. 17:368–376.

Gelfand AE, Smith AFM. 1990. Sampling-Based Approaches to Calculating Marginal Densities. Journal of the American Statistical Association. 85:398–409.

Gelman A, Rubin DB. 1992. Inference from Iterative Simulation Using Multiple Sequences. Statistical Science. 7:457–511.

Gilks WR, Richardson S, Spiegelhalter DJ. 1996. Markov Chain Monte Carlo in Practice, volume 39. Chapman & Hall/CRC.

Golubchik T, Brueggemann AB, Street T, Gertz RE, Spencer C, Ho T, Giannoulatou E, Link-Gelles R, Harding RM, Beall B, et al. (15 co-authors). 2012. Pneumococcal genome sequencing tracks a vaccine escape variant formed through a multi-fragment recombination event. Nature genetics. 44:352–5.

Grad YH, Kirkcaldy RD, Trees D, Dordel J, Harris SR, Goldstein E, Weinstock H, Parkhill J, Hanage WP, Bentley SD, et al. (11 co-authors). 2014. Genomic epidemiology of Neisseria gonorrhoeae with reduced susceptibility to cefixime in the USA: A retrospective observational study. The Lancet infectious diseases. 14:220–26.

Griffiths RC, Marjoram P. 1997. An ancestral recombination graph. Progress in population genetics and human evolution (Minneapolis, MN, 1994). 87:257–270.

Hamilton HL, Dillard JP. 2006. Natural transformation of Neisseria gonorrhoeae : From DNA donation to homologous recombination. Molecular Microbiology. 59:376–385.

Hanage W, Fraser C, Tang J, Connor T, Corander J. 2009. Hyper-Recombination, Diversity, and Antibiotic Resistance in Pneumococcus. Science. 324:1454–1457.

Harris SR, Feil EJ, Holden MT, Quail MA, Nickerson EK, Chantratita N, Gardete S, Tavares A, Day N, Lindsay JA, et al. (15 co-authors). 2010. Evolution of MRSA During Hospital Transmission and Intercontinental Spread. Science. 327:469–474.

Harrison OB, Maiden MC, Rokbi B. 2008. Distribution of transferrin binding protein B gene (tbpB) variants among Neisseriaspecies. BMC Microbiology. 8:66.

Hastie DI, Green PJ. 2012. Model Choice using Reversible Jump Markov Chain Monte Carlo. Statistica Neerlandica. 66:309–338.

Hedge J, Wilson DJ. 2014. Bacterial Phylogenetic Reconstruction from Whole Genomes Is Robust to Recombination but Demographic Inference Is Not. mBio. 5:e02158–14.

Höhna S, Drummond AJ. 2012. Guided Tree Topology Proposals for Bayesian Phylogenetic Inference. Systematic Biology. 61:1–11.

Holden MTG, Lindsay JA, Corton C, Quail MA, Cockfield JD, Pathak S, Batra R, Parkhill J, Bentley SD, Edgeworth JD. 2010. Genome sequence of a recently emerged, highly transmissible, multi-antibiotic- and antiseptic-resistant variant of methicillin-resistant Staphylococcus aureus, sequence type 239 (TW). Journal of bacteriology. 192:888–92.

Hudson RR. 1983. Properties of a neutral allele model with intragenic recombination. Theoretical Population Biology. 23:183–201.

Jukes T, Cantor C. 1969. Evolution of protein molecules. In: Mammalian Protein Metabolism, Academic Press, pp. 21–132.

Kennemann L, Didelot X, Aebischer T, Kuhn S, Drescher B, Droege M, Reinhardt R, Correa P, Meyer TF, Josenhans C, et al. (12 co-authors). 2011. Helicobacter pylori genome evolution during human infection. Proc Natl Acad Sci USA. 108:5033–5038.

Knorr-Held L, Richardson S. 2003. A Hierarchical Model for Space–Time Surveillance Data on Meningococcal Disease Incidence. Journal of the Royal Statistical Society Series C: Applied Statistics. 52:169–183.

Krebes J, Didelot X, Kennemann L, Suerbaum S. 2014. Bidirectional genomic exchange between Helicobacter pylori strains from a family in Coventry, United Kingdom. International Journal of Medical Microbiology. 304:1135–1146.

Kuhner MK, Felsenstein J. 1994. A simulation comparison of phylogeny algorithms under equal and unequal evolutionary rates. Molecular biology and evolution. 11:459–68.

Lartillot N, Philippe H. 2006. Computing Bayes Factors Using Thermodynamic Integration. Systematic Biology. 55:195–207.

Lin M, Kussell E. 2017. Correlated Mutations and Homologous Recombination Within Bacterial Populations. Genetics. 205:891–917.

Maiden MC, Bygraves JA, Feil EJ, Morelli G, Russell JE, Urwin R, Zhang Q, Zhou J, Zurth K, Caugant DA, et al. (13 co-authors). 1998. Multilocus sequence typing: A portable approach to the identification of clones within populations of pathogenic microorganisms. Proc Natl Acad Sci USA. 95:3140–5.

Manoharan-Basil SS, Gestels Z, Abdellati S, Akomoneh EA, Kenyon C. 2023. Evidence of horizontal gene transfer within porB in 19 018 whole-genome Neisseria spp. isolates: A global phylogenetic analysis. Microbial Genomics. 9:001041.

Mourkas E, Yahara K, Bayliss SC, Calland JK, Johansson H, Mageiros L, Muñoz-Ramirez ZY, Futcher G, Méric G, Hitchings MD, et al. (18 co-authors). 2022. Host ecology regulates interspecies recombination in bacteria of the genus Campylobacter. eLife. 11:e73552.

Nielsen R, Vaughn AH, Deng Y. 2025. Inference and applications of ancestral recombination graphs. Nature Reviews Genetics. 26:47–58.

Ozer EA, Nnah E, Didelot X, Whitaker RJ, Hauser AR. 2019. The population structure of Pseudomonas aeruginosa is characterized by genetic isolation of exoU+ and exoS+ lineages. Genome Biology and Evolution. 11:1780–1796.

Paradis E, Schliep K. 2019. Ape 5.0: An environment for modern phylogenetics and evolutionary analyses in R. Bioinformatics. 35:526–528.

Plummer M, Best N, Cowles K, Vines K. 2006. CODA: Convergence diagnosis and output analysis for MCMC. R News. 6:7–11.

Pupko T, Pe’er I, Shamir R, Graur D. 2000. A fast algorithm for joint reconstruction of ancestral amino acid sequences. Molecular Biology and Evolution. 17:890–6.

Rabiner L. 1989. A tutorial on hidden Markov models and selected applications in speech recognition. Proceedings of the IEEE. 77:257–286.

Robinson DF, Foulds LR. 1981. Comparison of phylogenetic trees. Mathematical Biosciences. 53:131–147.

Roijers F, Mandjes M, Van Den Berg H. 2007. Analysis of congestion periods of an m/m/-queue. Performance Evaluation. 64:737–754.

Saitou N, Nei M. 1987. The neighbor-joining method: A new method for reconstructing phylogenetic trees. Molecular biology and evolution. 4:406–25.

Sakoparnig T, Field C, Van Nimwegen E. 2021. Whole genome phylogenies reflect the distributions of recombination rates for many bacterial species. eLife. 10:e65366.

Schierup MH, Hein J. 2000. Consequences of recombination on traditional phylogenetic analysis. Genetics. 156:879–91.

Schliep KP. 2011. Phangorn: Phylogenetic analysis in R. Bioinformatics. 27:592–593.

Sheppard SK, Didelot X, Jolley KA, Darling AE, Pascoe B, Meric G, Kelly DJ, Cody A, Colles FM, Strachan NJC, et al. (24 co-authors). 2013. Progressive genome-wide introgression in agricultural Campylobacter coli. Molecular Ecology. 22:1051–1064.

Sheppard SK, Guttman DS, Fitzgerald JR. 2018. Population genomics of bacterial host adaptation. Nature Reviews Genetics. 19:549–565.

Spencer SE. 2021. Accelerating adaptation in the adaptive Metropolis–Hastings random walk algorithm. Australian and New Zealand Journal of Statistics. 63:468–484.

Stamatakis A. 2014. RAxML version 8: A tool for phylogenetic analysis and post-analysis of large phylogenies. Bioinformatics. 30:1312–1313.

Talts S, Betancourt M, Simpson D, Vehtari A, Gelman A. 2020. Validating Bayesian Inference Algorithms with Simulation-Based Calibration. arXiv. p. 10.48550/arXiv.1804.06788.

Vos M, Didelot X. 2009. A comparison of homologous recombination rates in bacteria and archaea. ISME Journal. 3:199–208.

Warren DL, Geneva AJ, Lanfear R. 2017. RWTY (R We There Yet): An R package for examining convergence of Bayesian phylogenetic analyses. Molecular Biology and Evolution. p. msw279.

Whidden C, Matsen FA. 2015. Quantifying MCMC Exploration of Phylogenetic Tree Space. Systematic Biology. 64:472–491.

Wilson IJ, Balding DJ. 1998. Genealogical inference from microsatellite data. Genetics. 150:499–510.

Xie W, Lewis PO, Fan Y, Kuo L, Chen MH. 2011. Improving Marginal Likelihood Estimation for Bayesian Phylogenetic Model Selection. Systematic Biology. 60:150–160.

Yahara K, Didelot X, Jolley KA, Kobayashi I, Maiden MCJ, Sheppard SK, Falush D. 2016. The landscape of realized homologous recombination in pathogenic bacteria. Molecular Biology and Evolution. 33:456–471.

Zhan SH, Wong Y, Ignatieva A, Eaton K, Guthrie I, Jeffery B, Palmer DS, Murall CL, Otto SP, Kelleher J. 2025. A Pandemic-Scale Ancestral Recombination Graph for SARS-CoV-2. bioRxiv. p. 10.1101/2023.06.08.544212.

Zhang C, Huelsenbeck JP, Ronquist F. 2020. Using Parsimony-Guided Tree Proposals to Accelerate Convergence in Bayesian Phylogenetic Inference. Systematic Biology. 69:1016–1032.

