## Supplementary Material for "Bayesian phylogenetic inference accounting for recombination in whole bacterial genomes"

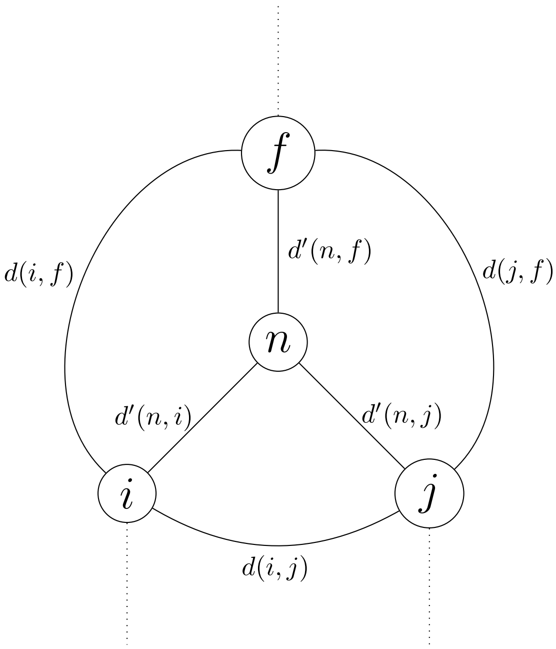


Figure S1: Illustration of how new branch lengths are proposed for a newly created node $n$ with parent $f$ and children $i$ and $j$.

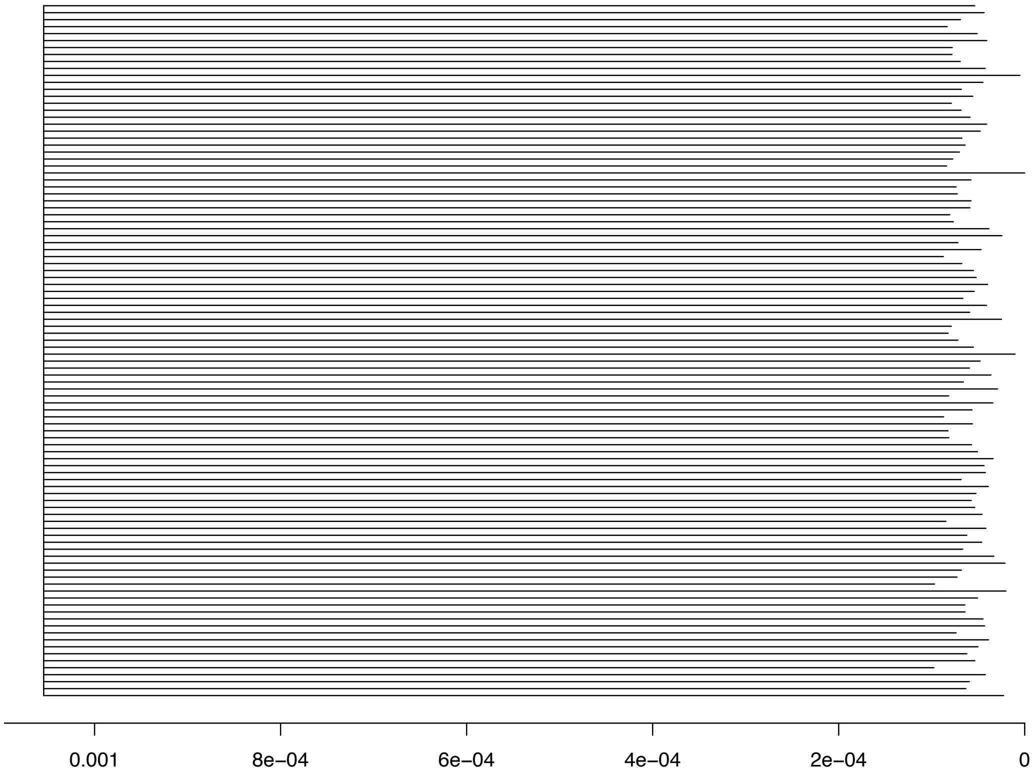


Figure S2: Star-like consensus phylogeny obtained when running without any data.
